# Altered Aperiodic EEG Activity in Adolescents with Neurofibromatosis Type 1

**DOI:** 10.64898/2026.09.28.754984

**Authors:** Marta Czime Litwińczuk, Thomas Donoghue, Jason Taylor, Gorana Pobric, Caroline Lea-Carnall, Shruti Garg

## Abstract

**Background:** The balance between excitatory and inhibitory neuronal signalling (E/I balance), proposed as a key mechanism controlling neuronal activity, is disrupted in many neurodevelopmental conditions. One such condition is Neurofibromatosis Type 1, a monogenic neurodevelopmental disorder characterised by increased inhibitory tone. While previous research suggests that E/I balance may be altered in infancy, it is unknown whether E/I balance remains altered during adolescence in NF1. Here we investigated this question using the aperiodic exponent of the resting electroencephalogram (EEG) power spectrum, a proposed electrophysiological index of E/I balance.

**Method:** The aperiodic exponent of the EEG power spectrum (‘1/f’) was estimated for 16 adolescents with NF1 diagnosis aged 11-16, and 16 age– and sex-matched neurotypical controls. General linear models (GLM) were fit to estimate effects of NF1 diagnosis, age and their interaction on aperiodic exponent. We also estimated the peak dominant frequency and power in oscillatory activity of the EEG signal and fitted the same structure of GLM models. Cognitive assessments across domains of memory, executive function and attention were also analysed with GLM to examine brain-cognition associations.

**Results:** The NF1 group had higher aperiodic exponent compared to neurotypical controls, but the aperiodic exponent was not associated with cognitive performance. After accounting for aperiodic activity, the NF1 group also had lower dominant oscillatory frequency than the control group. There was no significant interaction between NF1 diagnosis and age on PAF. There were no group differences in peak alpha power.

**Conclusions:** We demonstrate for the first time that the alterations to aperiodic activity in NF1 are consistent with the proposed mechanism of increased inhibitory tone. These findings show the potential of this paradigm for future pharmacological and brain stimulation trials investigating treatment-related changes in cortical function.

## Introduction

Neurofibromatosis Type 1 (NF1) is a monogenic, neurodevelopmental disorder with an estimated birth incidence of 1 in 2,700 (Evans et al., 2010). It is caused by mutation of the NF1 gene on chromosome 17q11.2, which encodes for the neurofibromin protein (Ballester et al., 1990). Clinical symptoms may include pigmentary lesions (café-au-lait spots), dermal neurofibromas, Lisch nodules, skeletal abnormalities, and optic gliomas (Ratner & Miller, 2015). The condition is also associated with difficulties in learning, and social and cognitive function, including domains of attention, executive function, cognitive flexibility, planning, working memory and inhibition (Gutmann et al., 2017; Hyman et al., 2005; Lehtonen et al., 2015). A substantial portion of individuals with NF1 also meet the criteria for diagnosis of attention deficit hyperactivity disorder (ADHD), autism spectrum conditions (ASC) and intellectual impairments (Garg et al., 2012; Gutmann et al., 2017; Hyman et al., 2005; Walsh et al., 2012).

Electroencephalography (EEG) records the brain’s electrical activity during rest and cognitive tasks and can be used to investigate the neural mechanisms of cognitive and behavioural difficulties in NF1. For example, Booth et al. (2023) investigated oscillatory dynamics during rest and a working memory task. During resting state, NF1 group showed increased delta (1-3 Hz) and theta (4-7 Hz) power, along with a lower peak alpha frequency (PAF). Further, while PAF showed a significant positive correlation with age in the control group, this relationship was absent in the NF1 group, suggesting a disrupted maturational trajectory of cortical oscillatory development. Booth et al. (2023) also analysed oscillatory activity during working memory, which revealed group differences in working memory task-related theta power and frontoparietal coherence. However, these group differences disappeared after adjusting task-related activity for resting-state activity, which suggests that the task-related differences reflected each group’s baseline level of activity.

Although Booth et al. (2023) reported differences in EEG activity in individuals with NF1, recent methodological advances have highlighted the importance of distinguishing oscillatory from aperiodic components of the EEG power spectrum (Donoghue et al., 2021). Aperiodic activity (with no characteristic frequency) contributes power across all frequencies; therefore, it can confound estimates of band specific power (Donoghue et al, 2020). Consequently, it remains unclear whether previously reported differences in EEG power and frequency in NF1 reflect genuine alterations in neural oscillations, changes in aperiodic activity, or a combination of both. To address this, we employ spectral parameterisation to separate oscillatory and aperiodic components of the EEG power spectrum, with two primary objectives: (i) to determine whether previously reported differences in spectral power and frequency persist when accounting for aperiodic activity, and (ii) to investigate whether NF1 is associated with alterations in aperiodic activity itself.

Aperiodic activity can be studied by measuring the aperiodic exponent from the power spectrum, whereby the exponent x of a 1/f^x fit to the data describes the rate of change in power across frequencies, characterising aperiodic activity. Gao et al. (2017) proposed that the aperiodic exponent may index the cortical excitation/inhibition (E/I) balance, because the time constants of the underlying excitatory (AMPA) and inhibitory (GABA) synaptic currents are thought to contribute to aperiodic activity (Alvarez & Destexhe, 2004; Pospischil et al., 2009). Specifically, computational modelling and animal research from Gao et al. (2017) predicts that greater inhibitory activity is associated with greater aperiodic exponent. Empirical work in humans shows systematic variation in aperiodic exponent across development (Stanyard et al., 2024), aging (Voytek et al., 2015) and clinical conditions (Donoghue, 2025), consistent with the exponent capturing meaningful individual and group differences in cortical activity.

The E/I balance is thought to be disrupted in NF1, as animal models show that the NF1 mutation leads to reduced or aberrant neurofibromin function with downstream increase in GABAergic activity (Costa et al., 2002). Additionally, animal and human pharmacological research supports the proposal that altered GABAergic activity underpins cognitive disruptions in NF1 (Bernardino et al., 2022; Cui et al., 2008; Mainberger et al., 2013). Neuroimaging studies provide further converging evidence for altered E/I balance, although these alterations appear region– and context-specific (Litwińczuk et al., 2026; Wild et al., 2026). Together, converging evidence across neurochemical, functional and computational neuroimaging approaches supports the hypothesis that NF1 is characterised by an altered E/I balance.

Motivated by the proposal that aperiodic exponent may also index E/I balance in NF1, Carter Leno et al. (2022) compared aperiodic exponent values between 10-month-old infants with NF1 and typically developing controls. During EEG data collection, infants passively watched videos. Infants with NF1 showed higher aperiodic exponent than the control group, which is consistent with the interpretation that individuals with NF1 exhibit a relatively increased inhibitory tone. Additionally, Carter Leno et al. investigated whether the aperiodic exponent could predict autistic traits at 36 months and whether executive attention would moderate this relationship. The results showed that higher aperiodic component values in infants with NF1 predicted greater autistic traits at 36 months, but only in infants with poorer attention at 24 months. This suggests that executive function may buffer the impact of early developmental E/I alterations in NF1. However, it is unknown whether the aperiodic exponent is also altered in NF1 during adolescence, a period during which neurochemical systems underlying E/I balance continue to mature (Caballero et al., 2021; Kilb, 2011; Perica et al., 2022). Neurodevelopmental conditions, such as ASC, ADHD and Fragile X syndrome, are characterised by alterations maturation of these systems (Bollmann et al., 2015; Port et al., 2016; Sohal & Rubenstein, 2019), suggesting that the maturation of mechanisms underlying E/I balance may also be altered in NF1, and such alterations may be detectable with aperiodic exponent.

To address these questions of putative difference in aperiodic activity in NF1, in this work, we investigate whether adolescents with NF1 diagnosis show altered EEG aperiodic exponent during rest, as compared to an age and sex matched neurotypical control group. We further explore aperiodic-controlled measures of oscillatory activity, and whether aperiodic exponent is related to assessments of executive function, attention and working memory performance. Following the findings reported in *Nf1* animal models, we hypothesise that the NF1 group, as compared to neurotypical controls, will show a higher aperiodic exponent consistent with increased inhibition. We further hypothesise that developmental characteristics of the aperiodic exponent and alpha wave characteristics will show reduced age-related changes in NF1, relative to controls.

## Methods

### Participants

This research presents a secondary analysis of the data presented by Pobric et al. (2021) and Booth et al. (2023). The data were collected as part of a pilot study [NCT03310996] designed to investigate effects and tolerability of non-invasive brain stimulation in NF1. The exclusion criteria were history of epilepsy, ongoing active treatment for any NF1 related complications (e.g. chemotherapy for optic glioma), or other clinically significant unrelated illness. Participants remained on their regular medication schedules. The study recruited 16 adolescents with an NF1 diagnosis aged 11-16 years and 16 age– and sex-matched typically developing controls.

Parents or guardians provided oral and written consent, and adolescents provided assent prior to participation. The control group had no pre-existing medical conditions or neurodevelopmental disorders. The NF1 group met the National Institute of Health’s NF1 diagnostic criteria (1988) with no history of epilepsy, no ongoing treatment for NF1-related complications or any other clinically significant unrelated illness. The NF1 participants were recruited through the Manchester Centre for Genomic Medicine, Neurofibromatosis charities, social media, and newsletters. Three of the NF1 participants were taking their regular medication schedule at the time of data collection.

### EEG acquisition and processing

EEG recordings were obtained using a ActiveTwo BioSemi system with 64 channels (positioned in standard 10-10 system), HEOG, VEOG and mastoids at sampling rate of 512Hz. During data collection, participants were asked to remain still and comfortable and avoid excessively moving their eyes or blinking. Participants completed 2.5 minutes of resting state with eyes-open, followed by 2.5 minutes of resting state with eyes closed. The session was followed by further cognitive tasks, described in full by Pobric et al. (2021) and Booth et al. (2023), but this study focuses only on the EEG recordings acquired during resting state.

Data processing was conducted in MatLab 2023a (The MathWorks, Inc., 2026), using EEGLab 2024.2 (Delorme & Makeig, 2004). The data were re-reference to linked mastoids, high– and low-pass filters were applied (0.1 Hz and 200 Hz, respectively), recordings were down-sampled to 256 Hz, a 48-52 Hz notch filter was applied, recordings were segmented to 2-second epochs without overlap between consecutive epochs and each epoch was mean-centred. Channel noise was assessed for channel removal using EEGLAB’s Clean Rawdata algorithm (with default values of flatline criterion set to 5 seconds, channel criterion was set to 0.8, and line noise criterion was set to 4), but no channels were removed. Artifact removal was performed with independent components analysis (ICA) generating 32 components and IClabel was called to automatically classify components into 8 classes (brain, eye, muscle, heart, line noise, channel noise, and other) (Pion-Tonachini et al., 2019). This classifier was pretrained by thousands of labelled components obtained through crowdsourcing and classifier training was done using scalp topography, power spectrum and autocorrelation between components. Components that were classified with ≥80% probability as artifact classes were removed, consistently with commonly used ICLabel-based preprocessing practices. This value was manually checked and compared to higher and lower thresholds (70-90%) on participant-by-participant basis to ensure that it was not removing neural sources. Rejected components themselves were also verified to ensure that neural signal sources were not being accidentally removed (Delorme, 2023). Finally, individual segments were inspected for short periods of noisy signal. Upon visual inspection, 3 NF1 and 3 control participants were rejected due to persistent noise in the data. In the remaining participants, in the control group the duration of recordings ranged between 102-210 seconds for eyes-open and 110-154 seconds for eyes-closed condition, and in the NF1 group duration of recordings ranged between 78-152 seconds for eyes-open and 66-148 seconds for eyes-closed condition. This discrepancy in recording duration was addressed in sensitivity analysis, described in later sections of this report.

### Spectral power parametrisation

After pre-processing, subsequent analyses were done in Python. First, power spectral density (PSD) was estimated for each epoch of the processed data using the multitaper method with discrete prolate spheroidal sequence (DPSS) tapers and a spectral smoothing bandwidth of 1 Hz, implemented in MNE-Python (v1.12.0) (Gramfort, 2013). PSDs were computed in the 2-30Hz frequency range with a frequency resolution of 0.5 Hz (derived from 2-s epochs) and averaged across epochs per channel.

The spectral parameterization method (specparam; formerly fooof, v1.1.1) (Donoghue et al., 2020) was applied to decompose power spectra into periodic components (oscillatory, rhythmic activity) and an aperiodic component. Spectral parameterization was applied using the standard aperiodic model (1/f, no knee), with peak width limits set to 1–6 Hz, a maximum of 6 peaks, a minimum peak height of 0.05, and a peak detection threshold of 1.5 standard deviations. Following recommendations for best practice (Ostlund et al., 2022), parameters were selected using a data-driven approach informed by the frequency range of interest and visual inspection of model fits in the full sample. Model fit was assessed using both explained variance (R²) and mean absolute error (MAE). At electrode POZ, model fit was good, with R² ranging from 0.973–0.995 for controls and 0.982–0.995 for the NF1 group during eyes-closed rest, and MAE ranging from 0.036-0.086 for controls and 0.040-0.077 for the NF1 group. We specifically focus the analyses on the electrode POZ, due to location of posterior dominant alpha rhythms (Freunberger et al., 2011; Klimesch, 1999; Mantini et al., 2007).

The aperiodic exponent was estimated from the aperiodic component, equivalent to the spectral slope in log-log space, with larger exponent values indicating a steeper power decay with increasing frequency. Additionally, dominant oscillatory power and frequency, controlling for aperiodic activity, were extracted from the 4-13 Hz frequency range. This band boundary was determined by identifying dominant peak frequencies estimated by the fitted spectral models for every participant, as recommended by Donoghue et al. (2020). In the control group dominant peak frequencies ranged 8.2–10.8 Hz, while in the NF1 group dominant peak frequencies ranged 4.9–12.3 Hz.

### Cognitive assessment measures

A battery of assessments was prepared to measure short-term memory, working memory, and attentional performance of the participants.

Short-term memory capacity was assessed using a computerised version of Corsi Blocks Task (Kessels et al., 2000) using PEBL software (Mueller & Piper, 2014). During this task, the participants view nine blue squares on the screen that change colour to orange (flash) for 500ms, with inter-stimulus-interval of 1000ms. The participants are asked to replicate the sequence by using the mouse. The task starts from 2 flashing blocks and increases in difficulty. The memory span is measured as maximum sequence that was correctly reproduced and the task was terminated when two sequences are incorrectly repeated.

Working memory capacity was assessed using Digit Span Forward and Digit Span Backward subtests of the Wechsler Intelligence Scale for Children – Fourth edition (WISC-IV) (Grizzle, 2011). During these tasks participants were asked to memorise a sequence of numbers and recall it either from the start-to-end or in reverse. Additionally, the participants completed adaptive visuospatial and auditory n-back tasks (Conway et al., 2005; Kane & Engle, 2002). Visuospatial task placed pictures of animals in one of 4 corners of the screen. The auditory task presented eight English consonants (c, g, h, k, p, q, t, w) spoken by a female voice. The visual and auditory stimuli were presented for 1000ms, with inter-stimulus-interval of 2500ms, with 14 non-target and 6 target stimuli. The participants were asked to respond when the current stimulus (image location or letter) matched the one presented N screens before (with N being 1, 2, or 3). The task was adaptive with difficulty of N increasing when the participant performed with ≥90% accuracy for a block or decreasing when the participant performed ≤70% accuracy. The task was terminated if performance stayed the same for two blocks. Performance was estimated as the average the average n-back level reached, and response time for target responses.

Attentional performance was measured with The Test of Everyday Attention for Children (TEA-Ch) (Manly et al., 2003), which includes the Sky Search, Score, Creature Counting, and Sky Search Dual Task. Raw scores for each subtest were adjusted for age based on standardised tables provided in the TEA-Ch manual.

### Cognitive data feature reduction

Feature reduction was performed on the digit span forward, digit span back, Corsi memory span, visuospatial and auditory N-back performance and mean response time during 2-back condition, and TEA-Ch dual task scores. Composite scores were created for the TEA-Ch battery: visual search efficiency was computed as average of Sky Search scores; executive attentional control was computed as average of Creature Counting scores.

All measures were standardised using z-score transformation and principal component analysis (PCA) was conducted. Components with eigenvalue ≥1.0 were retained for further analysis. Varimax rotation was applied to the retained component loadings, and component scores were estimated using the rotated loadings.

### General Linear Modelling

Shapiro-Wilk normality test showed normal distribution of the aperiodic exponent (controls W = .975, *p* = .948; NF1 W = .917, *p* = .220) and dominant oscillatory power (controls W = .907, *p* = .167; NF1 W = . 947, *p* = . 554), but not frequency (controls W = .888, *p* = . 090; NF1 W = . 860, *p* = . 038).

General linear models (GLM) were fitted to investigate the association between the aperiodic exponent at electrode POZ and NF1 diagnosis. Diagnosis was coded as a binary predictor (1 = NF1, 0 = control), age was centred through z-score transformation prior to analysis. The model included main effects of diagnosis and age, and age-diagnosis interaction. The same GLM structure was also separately fitted to dominant oscillatory power and frequency.

Since dominant oscillatory frequency showed non-normal distribution, permutation testing was implemented to assess significance of predictions in the dominant oscillatory frequency GLM model using the Freedman-Lane procedure (5,000 permutations), where for each predictor, a reduced model excluding that predictor was fitted, and its residuals were permuted and added back to the fitted values to generate null datasets. The full GLM was refit to each permuted dataset, and permutation-based p-values were computed as the proportion of permuted t-statistics exceeding the observed values.

To investigate associations between cognitive performance and aperiodic neural activity, GLMs were fit to PCA-derived scores. Predictors included the POZ aperiodic exponent (mean-centred), diagnosis (binary), age (z-scored) and the exponent-diagnosis interaction. This allowed investigation of group differences in associations between cognition and neural measures.

### Sensitivity analysis

We observed a discrepancy in duration of EEG recordings between groups, which could lead to cleaner, more stable estimates of PSD and aperiodic exponent in the control group. Therefore, we conducted an additional sensitivity analysis to investigate whether group differences in recording duration for the eyes-closed condition influenced the observed group effect. Specifically, we randomly subsampled the duration of control-group recordings during the eyes-closed condition to match the range observed in the NF1 group. For each control participant, we randomly selected a number of epochs drawn uniformly from the NF1 group’s range (33-74 epochs), then we randomly selected that number of epochs from the control participant’s complete data without replacement, and we re-computed the aperiodic exponent. The GLM was then re-fitted using the untouched NF1 group data and the down-sampled control-group data, and the resulting model parameters (β, t-statistic, and p-value) were recorded. This process was repeated 1000 times, with a random subset of epochs selected for the controls.

## Results

The following sections all report the results of analysis of eyes-closed condition. Figure 1 illustrates the fit of model at electrode POZ for each group and group differences, while Supplementary Material 1 presents model fits for individual participants. Aperiodic exponent findings were replicated during the eyes-open condition, and all models for eyes-open condition are presented in Supplementary Material 2.

**Figure 1.**
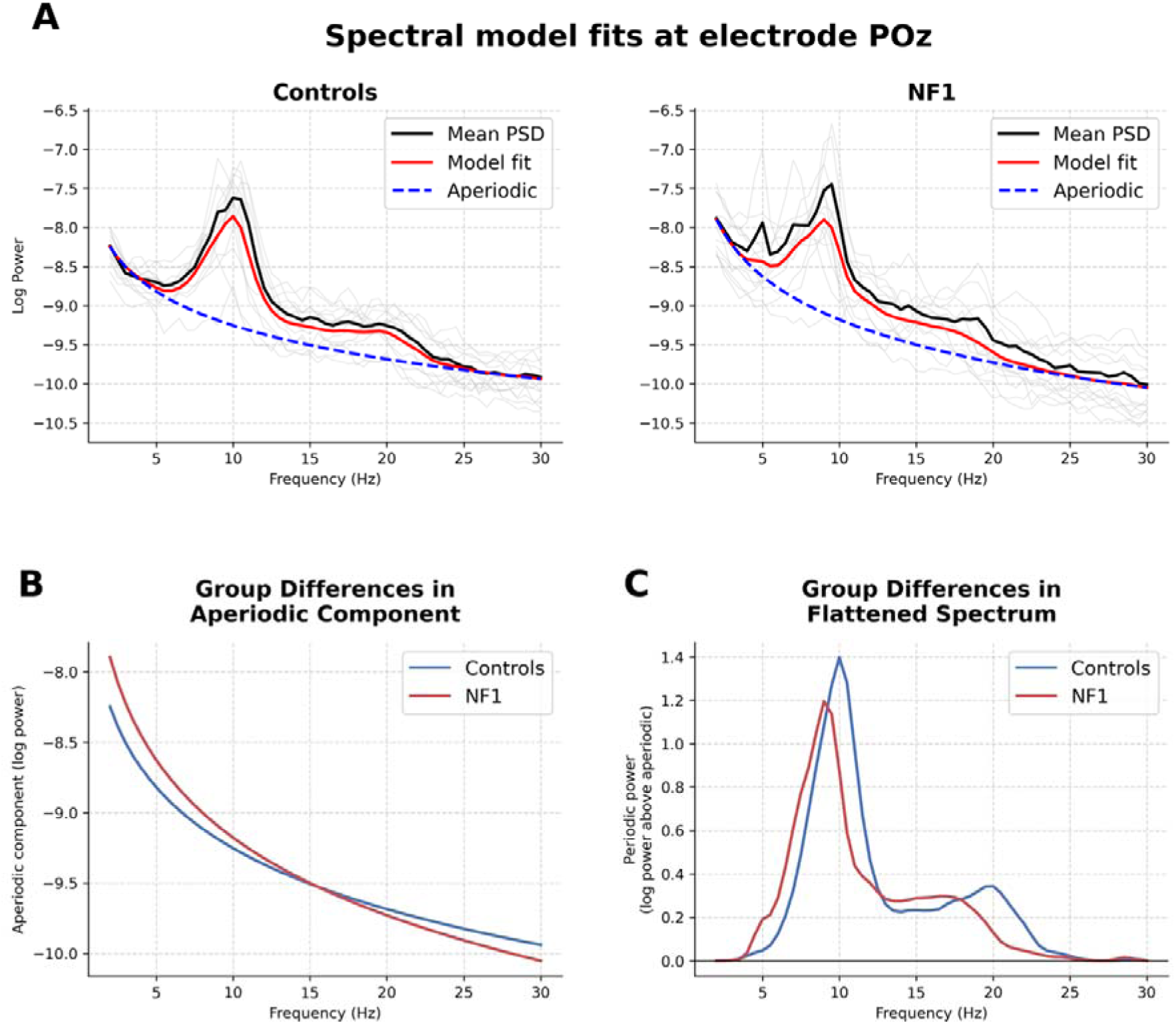
Spectral model fits at electrode POz during resting-state eyes-closed condition. (A) The group mean power spectra (PSD) and the spectral model fits. Pale grey lines represent individual participant power spectra. The black line represents the group mean power spectrum. The red line represents the mean of the spectral model fits estimated separately for each individual participant, and the blue dashed line represents the corresponding mean of individual aperiodic components. (B) Mean aperiodic components derived from individual spectral model fits for the control and NF1 groups, illustrating between-group differences in the broadband 1/f-like spectral component. (C) Mean periodic components (flattened spectra) obtained by subtracting the individual aperiodic component from the full spectral model fit, illustrating group differences in oscillatory activity after the aperiodic component was subtracted from PSD.

### Aperiodic exponent

Figure 2 presents the scalp distribution of the aperiodic exponent (A) and provides the boxplots for each group’s aperiodic exponent values at electrode POZ (B). The controls had mean exponent value of 1.437 (SD = 0.226), and NF1 had mean exponent value of 1.832 (SD = 0.330). The overall GLM model was significant, F(3, 22) = 4.617, *p* = .0119, explaining 38.6% of variance (R² = .386; adjusted R² = .303). The model showed a significant main effect of diagnosis, with the NF1 group showing a higher aperiodic exponent compared to the control group (β = 0.39, SE = 0.11, *t*(22) = 3.49, *p* = .002) (Figure 2B). There was no significant effect of age (β = –0.10, SE = 0.09, *t*(22) = –1.08, *p* = .292) or interaction between diagnosis and age (β = 0.14, SE = 0.12, *t*(22) = 1.21, *p* = .241).

**Figure 2.**
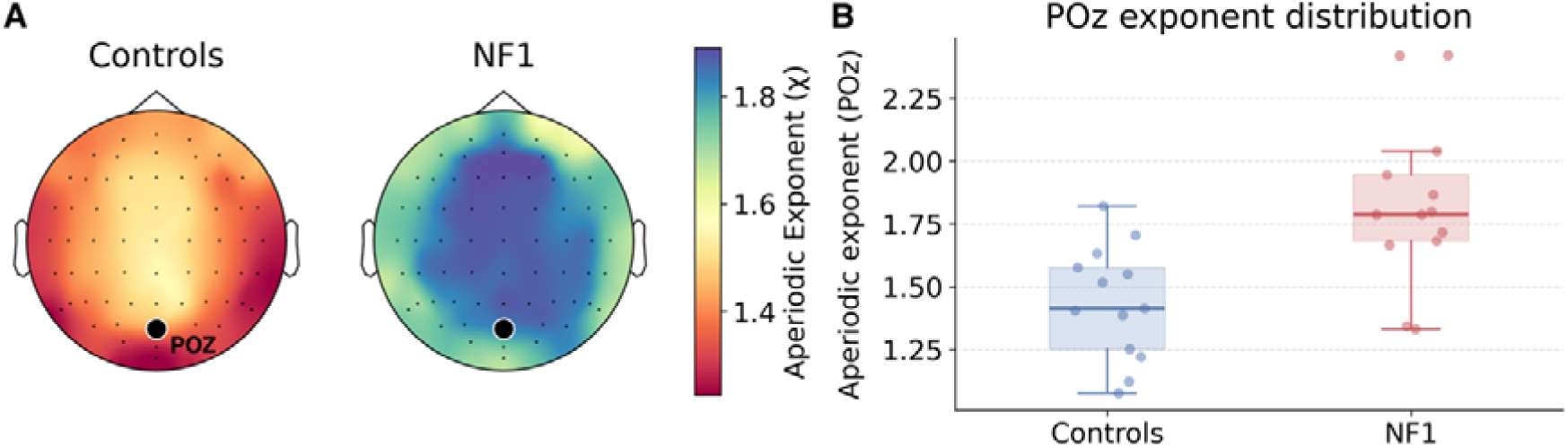
(A) The topological distribution of mean aperiodic exponent values estimated for the control and NF1 groups. (B) The aperiodic exponent values at electrode POZ for the control and NF1 group. The boxplots illustrate the median (line), interquartile range (box), and range within and range within 1.5× the interquartile range (whiskers). Individual points represent individual exponent values of every participant.

The sensitivity analysis revealed that the effect of diagnosis on aperiodic exponent was stable and not affected by the duration of recordings (mean β = 0.393, SD = 0.007; mean SE = 0.113; mean *t* = 3.47), with all 1000 resamples producing a significant effect of diagnosis (mean *p* = .002).

### Dominant oscillatory power and frequency

The estimated effects of NF1 diagnosis, age and interaction between diagnosis and age on dominant oscillatory power and frequency are illustrated in Figure 3. The dominant oscillatory power was 1.54 (SD = 0.36) in the control group, and 1.37 (SD = 0.38) in the NF1 group. However, the GLM model revealed no significant effects of diagnosis, age or their interaction.

**Figure 3.**
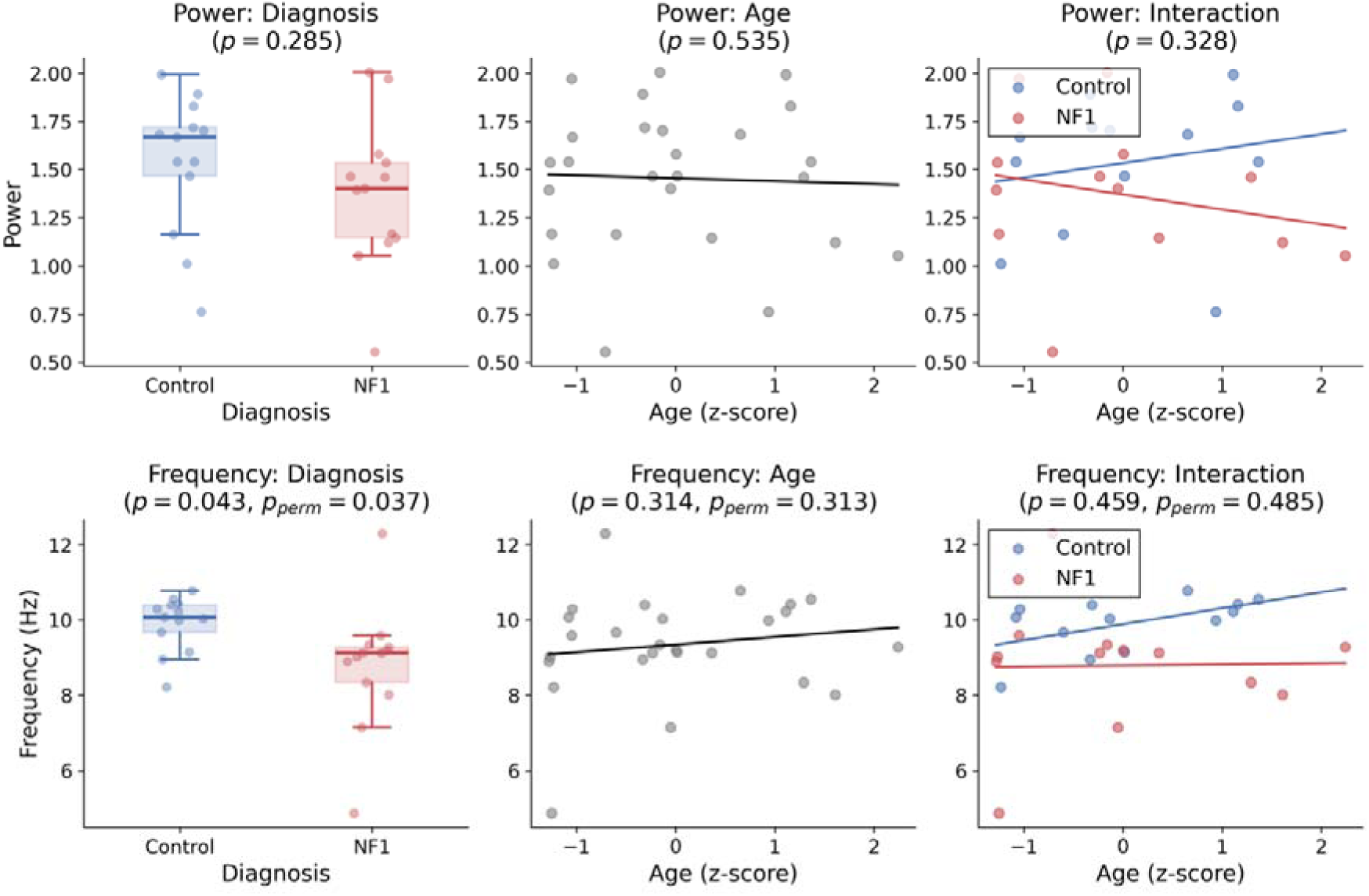
The scatter plots illustrate the GLM-derived effects of diagnosis, age and their interaction on oscillatory activity measures (power and frequency). Only effect of diagnosis on dominant oscillatory frequency reached statistical significance. The probability values are provided in panel titles (p = probability value, p_perm_ = permutation-based probability value).

The dominant oscillatory frequency at electrode POz was computed within the 4–13 Hz range. The NF1 group showed lower mean frequency (M = 8.79 Hz, SD = 1.65) than the control group (M = 9.90 Hz, SD = 0.73). The overall GLM model was not significant, F(3, 22) = 1.94, *p* = .153, explaining 20.9% of the variance (R² = .21; adjusted R² = .10). Within the model, there was a significant effect of diagnosis, where the NF1 group showed lower dominant oscillatory frequency than controls (β = –1.10, SE = 0.51, t(22) = –2.14, *p* = .043; *p_perm_* = .037). There were no significant effects of age or interaction between diagnosis and age.

### Cognition

Table 1 provides the descriptive statistics of each group’s performance on each cognitive measure.

**Table 1.** Mean and standard deviation, in brackets, of each group’s performance for each cognitive measure.

| Measure | Controls | NF1 |
| --- | --- | --- |
| Visuospatial N-back<br>(performance) | 2.34 (0.23) | 1.87 (0.28) |
| Auditory N-back (performance) | 2.68 (0.4) | 1.96 (0.37) |
| Visuospatial 2-back (response time) | 931.39 (319.33) | 965.9 (275.5) |
| Auditory 2-back (response time) | 1008.57 (197.53) | 1205.71 (323.93) |
| Corsi (memory span) | 5.58 (1.06) | 4.65 (1.05) |
| Digit span (forward) | 9.85 (1.72) | 7.62 (2.5) |
| Digit span (backward) | 8.62 (1.56) | 6.31(1.44) |
| Teach (Visual search) | 10.56 (1.78) | 9.54 (1.75) |
| Teach (Executive control) | 10.69 (2.55) | 8.65 (2.77) |
| Teach (Dual task) | 8.69 (1.44) | 8 (1.68) |

Three components were retained from principal component analysis, based on the Kaiser criterion (eigenvalues ≥ 1). These components accounted for 69.72% variance in the data (component 1 = 46.11%, component 2 = 12.48%, component 3 = 11.14%).

Following varimax rotation, the rotated component loadings were examined (Supplementary Material 3), which revealed that component 1 reflected working memory performance, component 2 reflected cognitive flexibility, and component 3 reflected speed-accuracy trade-off.

### Brain-cognition associations

GLM models were fitted to every component with diagnosis, age, aperiodic exponent at electrode POZ, and the interaction between diagnosis and aperiodic exponent as predictors.

For Component 1, reflecting working memory, the overall GLM model was significant, F(4, 21) = 8.35, *p* < .001, explaining 61.4% of variance (R² = .61; adjusted R² = .54). There was a significant effect of diagnosis, with NF1 participants showing lower component scores compared to controls (β = –2.96, SE = 0.71, *t*(21) = –4.18, *p* < .001). There was also a significant effect of age, with older participants showing higher scores (β = 0.80, SE = 0.29, *t*(21) = 2.75, *p* = .012). Neither the aperiodic exponent, nor the interaction between diagnosis and exponent were significant.

The GLM models of Components 2 and 3 did not reach statistical significance and revealed no significant effects of diagnosis, age, aperiodic exponent, or their interactions.

## Discussion

Convergent animal, pharmacological and neuroimaging evidence points towards altered E/I balance as a key physiological mechanism of NF1 (Carter Leno et al., 2022; Costa et al., 2002; Cui et al., 2008). In this study, we investigated whether adolescents with NF1 show altered aperiodic exponent in EEG data, which has been proposed to index E/I balance (Gao et al., 2017). The NF1 group showed higher aperiodic exponent during rest compared to neurotypical controls. However, this group difference in aperiodic exponent was not related to cognitive performance. Additionally, the NF1 group had lower dominant oscillatory frequency than the control group, after accounting for aperiodic activity. Further, there were no group differences in dominant oscillatory power. Together these findings demonstrate neurophysiological differences in adolescents with NF1, consistent with altered E/I balance.

Our hypothesis that NF1 would be characterised by greater inhibitory activity, reflected by greater aperiodic exponent, was motivated by converging genetic animal models of NF1, and pharmacological treatment studies and human research in NF1 (Bernardino et al., 2022; Carter Leno et al., 2022; Costa et al., 2002; Cui et al., 2008; Mainberger et al., 2013; Pobric et al., 2021). Our results provide further evidence supporting this hypothesis, as adolescents with NF1 showed a higher aperiodic exponent during rest than neurotypical controls. This is consistent with previous findings in infants (Carter Leno et al., 2022), and suggests that the altered aperiodic exponent may represent a stable neurophysiological signature of NF1 across development (Petri et al., 2025). Further, the aperiodic exponent is one proposed index of the E/I balance, where greater inhibitory tone is associated with higher aperiodic exponent (Gao et al., 2017). Therefore, our finding that NF1 adolescents show higher aperiodic exponent is consistent with the hypothesis of individuals with NF1 have a relatively elevated inhibitory tone, as predicted by animal models of GABAergic activity in NF1 (Costa et al., 2002).

Our findings provide a complementary electrophysiological perspective on E/I balance in NF1, extending previous research using fMRI, MRS and PET (Doherty et al., 2023; Mainberger et al., 2013; Ribeiro et al., 2015; Violante et al., 2016; Violante et al., 2013). However, the wider neuroimaging literature suggests that E/I alterations in NF1 are complex, varying across brain regions and cognitive contexts. A recent systematic review from our research group found little support for a global increase in inhibition in NF1(Wild et al., 2026). Previous MRS and PET studies have reported reduced GABA availability and decreased GABA-A receptor binding in specific cortical and subcortical regions (Violante et al., 2016; Violante et al., 2013). Although these findings appear inconsistent with our interpretation of increased inhibitory activity, we note that the different methodologies capture distinct aspects of neurophysiology. Further evidence from fMRI suggests that NF1 may be associated with altered modulation of neural activity in response to cognitive demands (Litwińczuk et al., 2026; Zang et al., 2012; Zimerman et al., 2015). Collectively, these findings suggest that NF1 may be characterised by region and context-dependent alterations in E/I balance, rather than a global shift towards increased inhibition. Specifically, in the context of our resting state EEG recordings from adolescents with NF1, our findings do point towards increased inhibitory tone, likely reflecting one aspect of a broader, region– and context-specific E/I profile.

We followed the investigation of diagnosis-related group differences in aperiodic exponent with investigation of developmental differences, focusing on the interaction between NF1 diagnosis and age. We expected that adolescents with NF1 would show altered age-related variation in aperiodic exponent, because research shows that biological mechanisms of E/I balance continue to mature during adolescence (Caballero et al., 2021; Kilb, 2011; Perica et al., 2022). However, we found no evidence for such differences, but future work should replicate this in larger sample sizes. Future research should therefore investigate the relationship between aperiodic activity and the neurotransmitter systems responsible for regulating E/I balance across development with a larger sample size.

While Booth et al. (2023) applied traditional analysis of spectral power to resting state EEG recordings from adolescents with NF1, we separated the oscillatory and aperiodic components within the 4-13 Hz range from the same dataset. Consistent with Booth et al. (2023), our analysis showed lower dominant oscillatory frequency in NF1 than in neurotypical controls. However, while Booth et al. (2023) reported a positive age-PAF correlation in neurotypical control group that was absent in the NF1 group, we did not find evidence for an interaction between age and diagnosis, although the pattern of coefficients was consistent. This difference between studies may emerge from differences in statistical approaches, our smaller sample size and separate analysis of eyes open and closed conditions, or genuine differences introduced by separating periodic and aperiodic activities.

Replication in a larger sample will be necessary to further clarify the relationship between resting state dominant oscillatory activity and age in NF1. Importantly, by applying spectral parameterization, we demonstrate that the previously observed reduction in PAF in NF1 (Booth et al., 2023) is not driven by differences in aperiodic activity but instead reflects altered dynamics of the oscillations themselves. The findings also differed in other aspects of oscillatory activity: Booth et al. (2023) reported group differences in delta and theta power, which were not examined as separate measures in the present analysis. However, our finding of a robust difference in aperiodic exponent suggests that it is plausible that these previously reported group differences in delta and theta power may partly reflect differences in aperiodic activity. Finally, neither our study nor Booth et al. (2023) found any group difference in oscillatory power.

Finally, we also investigated whether baseline aperiodic activity was related to cognitive performance across domains of executive function, attention and memory. Factor analysis resulted with three components, that appeared to reflect Working Memory (characterised by high loading of N-back performance, Digit span backward), Attention Control (characterised by high loading on TEA-Ch dual task performance and negative loading on auditory 2-back response time), and Speed– accuracy Trade-off (characterised by longer visuospatial 2-back response times, modest positive loading on visuospatial 2-back performance, and negative loading on Corsi blocks memory span). While NF1 diagnosis and age both affected participant scores on working memory factor, there was no effect of the aperiodic exponent or interaction between aperiodic exponent and NF1 diagnosis. Together, these analyses show that there is no evidence to suggest that aperiodic activity during resting state directly relates to cognitive performance across these domains. Cognitive performance likely depends more strongly on neural dynamics during tasks than baseline resting-state activity alone. Indeed previous research has shown that task-related changes in aperiodic activity are related to working memory (Virtue-Griffiths et al., 2025), attention (Waschke et al., 2021) and cognitive processing speed (Ouyang et al., 2020), suggesting that modulation of the aperiodic exponent during task performance may relate to cognitive ability in NF1.

### Limitations

Importantly, the aperiodic exponent is not a direct measure of E/I balance. Although it can reflect E/I balance under specific assumptions (Gao et al., 2017), similar 1/f-like dynamics can arise from other mechanisms, including recurrent network activity, neuronal synchrony and population spiking statistics (Donoghue, 2025; Preston et al., 2026). Therefore, while our work provides convergent evidence with other research for elevated inhibitory activity in NF1 during rest, interpretation of the aperiodic exponent should be approached with caution, and it should not be considered a direct measure of GABAergic function.

Further, this study analysed a cross-sectional dataset, while modelling of longitudinal data could accurately characterise developmental trajectories and allow for developmental prognosis. Longitudinal data could also reduce effects of NF1 heterogeneity (e.g. severity, genetic variants) on statistical power to detect neurophysiological alterations to brain development. As result we can only discuss how electrophysiological markers vary across age, and we cannot establish how these markers change with age. The small sample size included in this study may have further reduced our sensitivity to detect interaction effects and associations in the presence of NF1 heterogeneity. Additionally, there are many sources of heterogeneity that we could not account for due to this small sample size. Meanwhile, some individuals with NF1 take medications to reduce the impact of their clinical symptoms, NF1 can emerge from a variety of genetic mutations, it may be inherited or de-novo mutation, its severity may vary across individuals, and it may affect different aspects of cognition (Pasmant et al., 2012). Importantly, some participants within this sample also had comorbid neurodevelopmental diagnoses, including ADHD and autism spectrum conditions (ASC) (Gutmann et al., 2017; Hyman et al., 2005; Walsh et al., 2012). Such comorbidities may contribute to electrophysiological differences, particularly given that aperiodic activity has been shown to vary across neurodevelopmental conditions, including ADHD and ASC, although findings are inconsistent across studies (Donoghue, 2025). Overall, these sources of heterogeneity should be considered when interpreting the observed electrophysiological differences.

### Conclusions

In conclusion, adolescents with NF1 show increased aperiodic exponent and reduced dominant oscillatory frequency relative to neurotypical controls. Together, these findings provide further evidence of altered neurophysiological dynamics in NF1 and are consistent with an alteration in E/I balance, potentially involving increased inhibitory influence during rest. Future longitudinal and task-based research will be important for clarifying how these alterations relate to developmental changes in E/I balance and cognitive function in NF1, and future research combining electrophysiological and neurochemical neuroimaging measures will be important for clarifying the development of GABAergic systems and their influence on neural dynamics.

## Data availability

The datasets used and/or analysed during the current study are available from the corresponding author on reasonable request.

## Ethics statement

Ethics approval for the study was obtained from the Greater Manchester West Research Ethics Committee (17/NW/0364). Written informed consent was obtained from the parents and older adolescent participants and assent was obtained from the younger participants.

