## Supplementary Materials for "Altered Aperiodic EEG Activity in Adolescents with Neurofibromatosis Type 1"

### Supplementary Material 1

Supplementary Figures 1 and 2 respectively present model fits for individual participants in control and NF1 group.

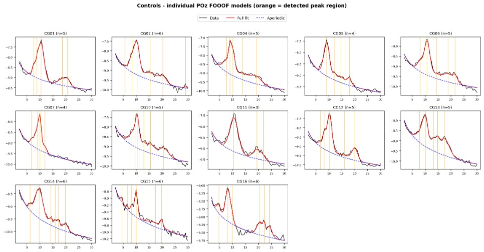

Supplementary Figure 1 The individual participant power spectra density (PSD) and the spectral model fits in the neurotypical control group. The black line represents the PSD. The red line represents the spectral model fit estimated separately for each individual participant, and the blue dashed line represents the corresponding individual aperiodic components. Orange lines represent detected peaks.

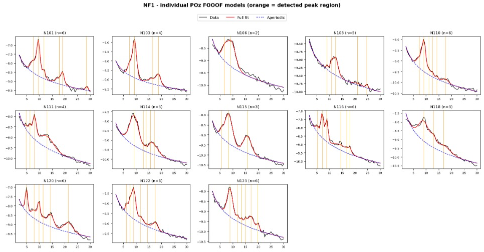

Supplementary Figure 2 The individual participant power spectra density (PSD) and the spectral model fits in the NF1 group. The black line represents the PSD. The red line represents the spectral model fit estimated separately for each individual participant, and the blue dashed line represents the corresponding individual aperiodic components. Orange lines represent detected peaks.

### Supplementary Material 2

Overview

The following supplementary results replicate the analysis presented in the main manuscript. While the analysis in the main manuscript were conducted on data obtained during eyes closed condition of resting state data, these analysis were conducted on data obtained during eyes open condition.

Diagnostics

Following processing of EEG data, the recoding duration ranged between 102-210 seconds for the control group, and 78-152 seconds for the NF1 group.

The model fit (R²) at electrode POZ ranged between 0.947 – 0.992 for the control group and 0.974 – 0.996 for the NF1 group. The mean absolute error (MAE) at electrode POZ ranged between 0.029 – 0.066 for the control group and 0.034 – 0.077 for the NF1 group.

Shapiro-Wilk normality test showed normal distribution of the aperiodic exponent (controls: W = 0.936, p = .408; NF1: W = 0.920, p = .248), but not the dominant oscillatory frequency (controls: W = 0.843, p = .023; NF1: W = 0.858, p = .036) or dominant oscillatory power (controls: W = 0.804, p = .008; NF1: W = 0.955, p = .672).

GLM of aperiodic exponent

The aperiodic exponent was 1.27 (SD = 0.19) in the control group and 1.62 (SD = 0.31) in the NF1 group. The overall GLM model of the aperiodic exponent at electrode POZ was significant, F(3, 22) = 5.23, p = .007, explaining 41.6% of variance (R² = .42; adjusted R² = .34). The model showed a significant main effect of diagnosis with the NF1 group showing a higher aperiodic exponent compared to control group (β = 0.34, SE = 0.10, t(22) = 3.46, p = .002). The effect of age was not significant (β = -0.14, SE = 0.08, t(22) = -1.81, p = .084). The interaction between age and diagnosis was not significant (β = 0.16, SE = 0.10, t(22) = 1.53, p = .139).

GLM of dominant oscillatory power

The dominant oscillatory power was 0.57 (SD = 0.50) in the control group and 0.89 (SD = 0.46) in the NF1 group. The overall GLM model of dominant oscillatory power was not significant, F(3, 22) = 0.86, p = .479, explaining 10.4% of the variance (R² = .10; adjusted R² = −.02). The model showed no significant main effect of diagnosis (β = 0.31, SE = 0.20, t(22) = 1.60, p = .124; pperm = .122), indicating that dominant oscillatory power did not differ significantly between the NF1 and control groups. The effect of age was also not significant (β = 0.004, SE = 0.16, t(22) = 0.02, p = .982; pperm = .980), nor was the interaction between age and diagnosis (β = 0.01, SE = 0.20, t(22) = 0.06, p = .952; pperm = .958).

GLM of dominant oscillatory frequency

The dominant oscillatory frequency was computed within the 4–13 Hz range. The control group showed a mean frequency of 10.14 Hz (SD = 1.46), while the NF1 group showed a mean of 8.10 Hz (SD = 1.76). The overall GLM model was significant, F(3, 22) = 4.35, p = .015, explaining 37.2% of the variance (R² = .37; adjusted R² = .29). The model showed a significant main effect of diagnosis, with the NF1 group showing lower dominant oscillatory frequency than controls (β = -2.00, SE = 0.63, t(22) = -3.19, p = .004; pperm = .004). The effect of age was not significant (β = 0.80, SE = 0.50, t(22) = 1.59, p = .126; pperm = .114). The interaction between age and diagnosis was not significant (β = -0.76, SE = 0.64, t(22) = -1.19, p = .248; pperm = .241).

### Supplementary Material 3

Supplementary Figure 4 presents the varimax the rotated component loadings of principal component analysis.

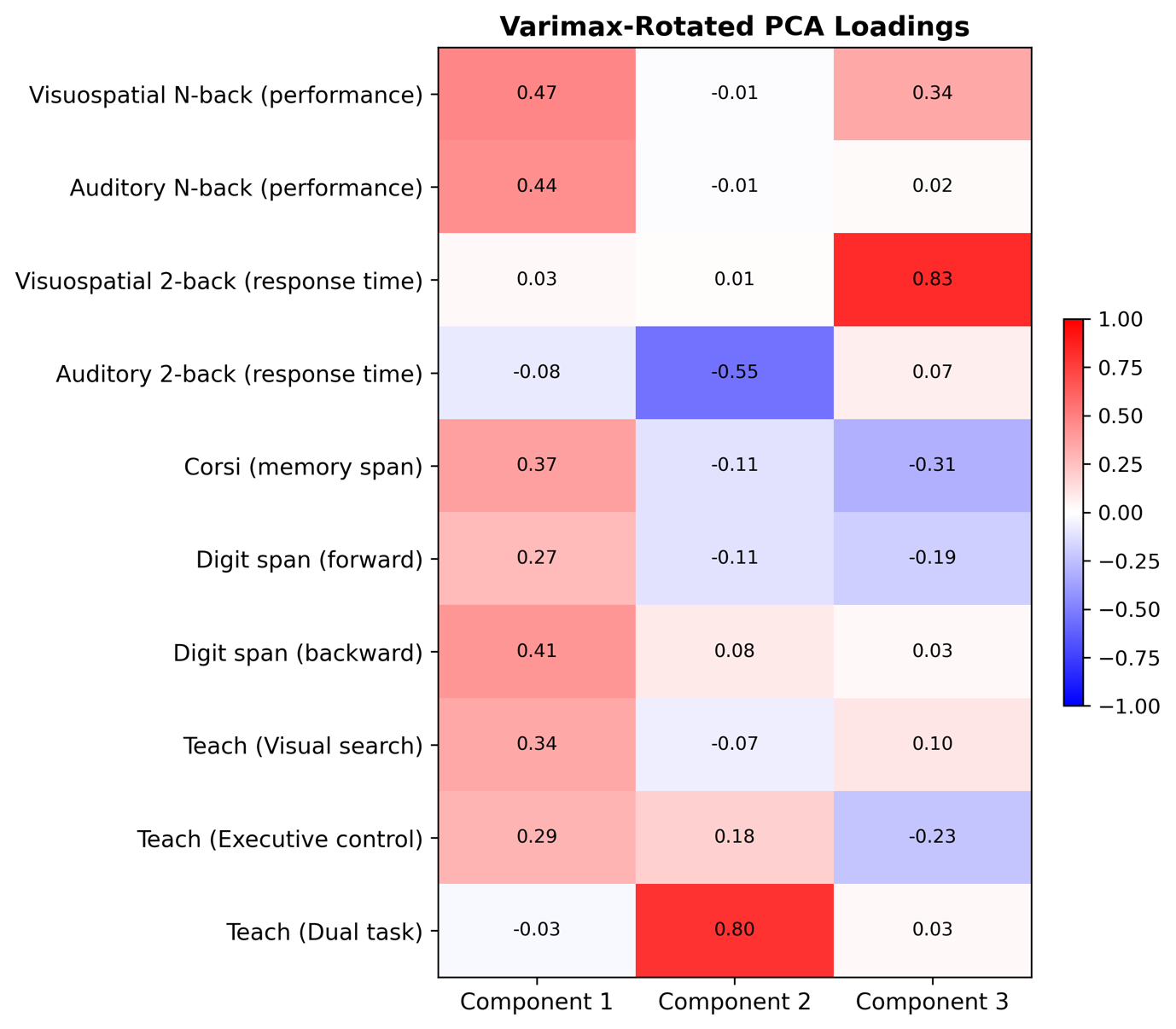

Supplementary Figure 3 The varimax-rotated PCA loadings for the 3 components that passed the Keiser criterion (eigenvalues ≥ 1).
